# The SARS-CoV-2 protein NSP2 impairs the microRNA-induced silencing capacity of human cells

**DOI:** 10.1101/2022.01.25.477753

**Authors:** Limei Zou, Clara Moch, Marc Graille, Clément Chapat

## Abstract

The coronavirus SARS-CoV-2 is the cause of the ongoing pandemic of COVID-19. Given the absence of effective treatments against SARS-CoV-2, there is an urgent need for a molecular understanding of how the virus influences the machineries of the host cell. The SARS-CoV-2 generates 16 Non-Structural Proteins (NSPs) through proteolytic cleavage of a large precursor protein. In the present study, we focused our attention on the SARS-CoV-2 protein NSP2, whose role in the viral pathogenicity is poorly understood. Recent proteomic studies shed light on the capacity of NSP2 to bind the 4EHP-GIGYF2 complex, a key factor involved in microRNA-mediated silencing of gene expression in human cells. In order to gain a better understanding of the function of NSP2, we attempted to identify the molecular basis of its interaction with 4EHP-GIGYF2. Our data demonstrate that NSP2 physically associates with the endogenous 4EHP-GIGYF2 complex in the cytoplasm. Using co-immunoprecipitation and *in vitro* interaction assays, we identified both 4EHP and a central segment in GIGYF2 as binding sites for NSP2. We also provide functional evidence that NSP2 impairs the function of GIGYF2 in mediating mRNA silencing using reporter-based assays, thus leading to a reduced activity of microRNAs. Altogether, these data reveal the profound impact of NSP2 on the post-transcriptional silencing of gene expression in human cells, pointing out 4EHP-GIGYF2 targeting as a possible strategy of SARS-CoV-2 to take over the silencing machinery and to suppress host defenses.

## Introduction

Beta-coronaviruses (β-CoVs) are enveloped RNA viruses that infect a variety of vertebrate hosts, including humans (1). In the last decades, two β-CoVs have caused epidemic diseases of the respiratory tract: severe acute respiratory syndrome (SARS-CoV-1) in 2002 (2,3) and Middle East respiratory syndrome (MERS) in 2012 (4). A new β-CoV (SARS-CoV-2) emerged in 2019 that is the causative agent of coronavirus disease 2019 (COVID-19) pandemic (5). These three viruses possess single-stranded, positive-sense RNA genomes of nearly 30 kb in length (6). The SARS-CoV-2 genome encodes 29 proteins with multiple functions in virus replication and packaging, including 4 structural proteins (the nucleocapsid N, envelope E, membrane M, and spike S proteins), 7 accessory proteins (ORF3a–ORF8) whose functions in SARS-Cov-2 pathogenesis remain largely unknown, and 16 non-structural proteins (NSP1–NSP16) that encode the RNA-directed RNA polymerase, helicase, protease, and other components required for virus replication (for review, see (7)).

Due to the urgent need to better understand SARS-CoV-2 biology, several CRISPR-Cas9 and proteomic-based screening campaigns investigated the landscape of host factors which are targeted by the virus proteome (8–12). These screens identified more than 300 high-confidence protein-protein interactions between human and SARS-CoV-2 proteins, highlighting the intimate connection of SARS-CoV-2 proteins with multiple biological processes, including protein trafficking, transcription and mRNA translation. Among these interactions, the Non-Structural Protein 2 (NSP2) has been found to interact with key host proteins involved in vesicle trafficking (FKBP15, WASHC) and mRNA translation (4EHP, GIGYF2) which could be of therapeutic importance (10,12).

NSP2 exists in all coronaviruses studied to date, including SARS-CoV-1, SARS-CoV-2, MERS, and in their closely related β-CoVs infecting mammals. Although its role in the SARS-CoV-2 pathogenicity has not been fully elucidated, the deletion of NSP2 in SARS-CoV-1 attenuates viral growth and RNA synthesis (13). Recent findings showed that SARS-CoV-2 NSP2 is undergoing positive nature selection and could be thus essential to the virus (14,15). At the structural level, NSP2 has a complex multi-domain topology including an N-terminal domain with a highly-conserved zinc binding site, and a C-terminal region rich in β-strands. With the exception of the zinc binding site, NSP2 displays a rapidly evolving surface with the presence of natural variations that could impact host-virus interactions (16–19). Among its host interactors, SARS-CoV-2 NSP2 interacts with the 4EHP-GIGYF2 complex, a key machinery in translational silencing and mRNA decay (10–12,20). It is worth noting that the NSP2/4EHP-GIGYF2 association is conserved across SARS-CoV-1 and MERS, corroborating a functional importance for β-CoV infection in general (11,20).

The cap-binding eIF<u>4E</u>–<u>H</u>omologous <u>P</u>rotein (4EHP) is an integral component of post-transcriptional silencing mechanisms through competing with the eIF4F complex for binding to the mRNA 5’cap (21). In complex with the <u>G</u>RB10-Interacting <u>GYF</u> (glycine-tyrosine-phenylalanine domain) protein 2 (GIGYF2), the cap-binding activity of 4EHP is required for the optimal translational repression by microRNAs (miRNAs), as well as the RNA-binding proteins ZNF598 and Tristetraprolin (TTP) (22–28). In the case of miRNA-driven silencing, the recruitment of 4EHP-GIGYF2 is initiated by the miRNA-induced silencing complex (miRISC), an assembly of argonaute and TNRC6/GW182 proteins. 4EHP can be physically mobilized through the interaction of the GYF domain of GIGYF2 with a proline-proline-glycine-leucine (PPGL) motif in TNRC6/GW182 (29,30). At the functional level, this 4EHP/miRNA axis is required to control ERK signaling, as well as to suppress IFN-β production by effecting the *miR-34a*-induced translational silencing of *Ifnb1* mRNA (31,32).

Beyond miRNA action, 4EHP-GIGYF2 also forms a translation inhibitory complex with the RNA-binding protein ZNF598, which functions in ribosome stalling on internally polyadenylated mRNAs during ribosome quality control (22,33). ZNF598 is also necessary for the repression of TTP-targeted mRNAs that encode inflammatory cytokines. The 4EHP-GIGYF2-ZNF598 complex binds TTP during an innate immune response in mouse macrophages to control the production of TTP-targeted mRNAs such as *TNF-α*, *Ier3*, *Csf2*, and *Cxcl10.* In all cases, TNRC6/GW182, ZNF598 and TTP display a comparable binding mode to GIGYF2, namely via the recognition of a proline stretch by the GYF domain of GIGYF2 (23,34).

A recent genetic screen has revealed that both 4EHP and GIGYF2 are necessary for infection by SARS-CoV-2 *in vitro*, while dispensable for seasonal coronaviruses (8). This contribution of 4EHP-GIGYF2 could likely originate from their interaction with NSP2, and thus serves as a potential drug target for anti-SARS-CoV-2 drug development. In the present report, we focused on the physical and functional interplays existing between the 4EHP-GIGYF2 complex and the SARS-CoV-2 protein NSP2 in human cells. Combining interaction assays and reporter-based approaches, our data shed light on the profound impact of NSP2 on the 4EHP-GIGYF2-mediated translational silencing of gene expression.

## Results

### NSP2 binds the 4EHP-GIGYF2 complex ***in cellulo***

Early large scale studies reported the capacity of SARS-CoV-2 NSP2 to bind the 4EHP-GIGYF2 complex using affinity-purification mass spectrometry (10,11). To validate the physical association of NSP2 with the GIGYF2-4EHP complex, we first sought to detect their interaction using co-immunoprecipitation (co-IP). An inducible Flag-tagged version of NSP2 was stably expressed in HEK293 Flp-In T-REX cells and co-IPs were performed following tetracycline-induced expression of Flag-NSP2. Lysates prepared from control, non-induced, and induced cells were immunoprecipitated with anti-Flag antibody. Subsequent Western blot (WB) analysis of the co-IP fraction showed that Flag-NSP2 efficiently binds both endogenous GIGYF2 and 4EHP (Fig. 1A). This interaction was similarly detected in RNase A-treated lysates, indicating that the NSP2/4EHP-GIGYF2 interaction occurs in an RNA-independent manner. The GIGYF2-associated protein ZNF598 was also found along with NSP2, while CNOT9, a subunit of the CCR4-NOT complex known to bind GYGYF2, was not detected (35) (Fig. 1A). Together, these data demonstrate that NSP2 physically associates with the 4EHP-GIGYF2 complex, as well as their interacting protein ZNF598.

**Figure 1.**
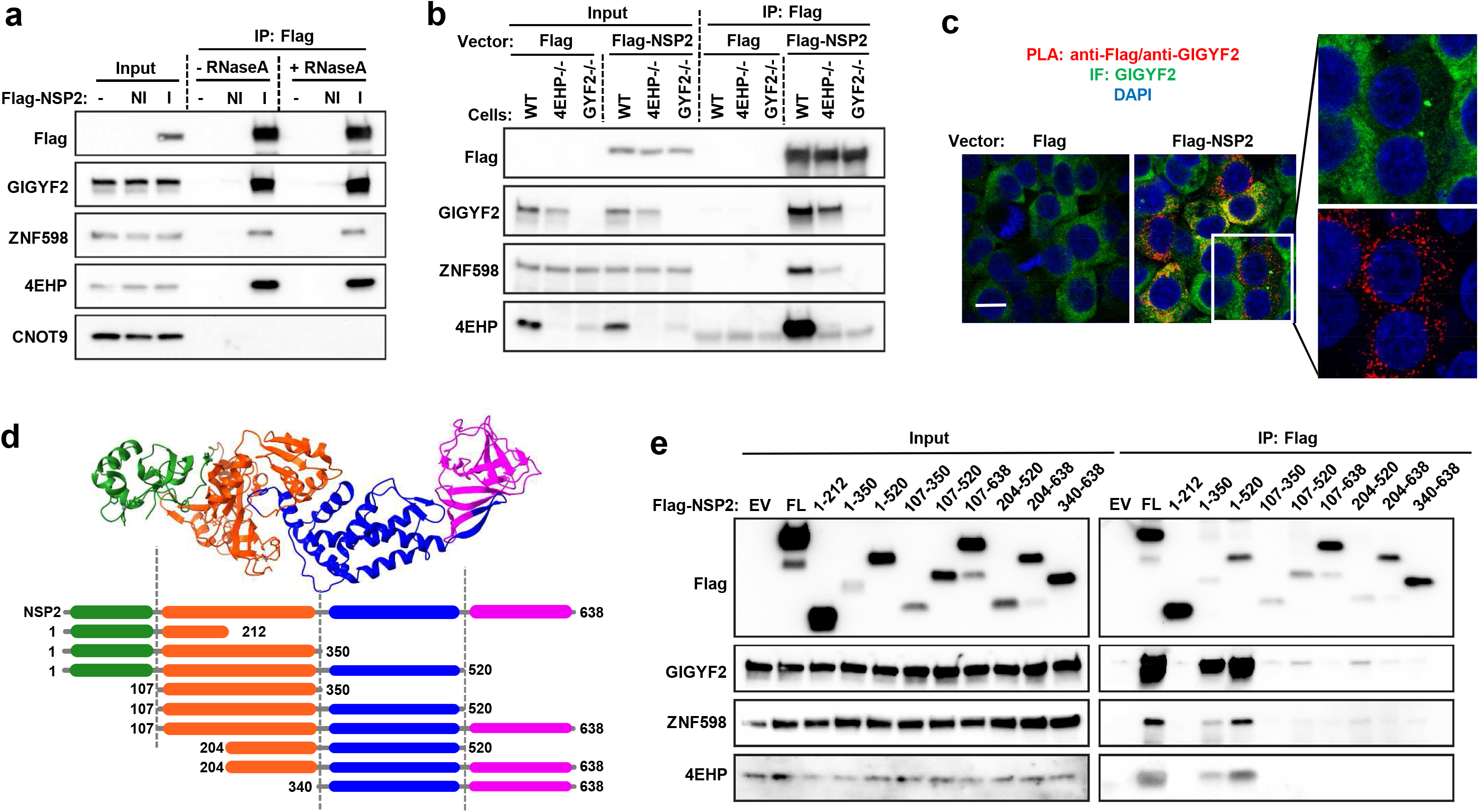
NSP2 interacts with the 4EHP-GIGYF2 complex in the cytoplasm. **(A)**Co-Immunoprecipitation (co-IP) between Flag-NSP2 and endogenous 4EHP-GIGYF2. A tetracycline-inducible Flag-NSP2 construct was stably transfected in HEK293 flp-In T-Rex. Extracts from untransfected (-), non-induced (NI) or tetracycline-induced (I) cells were immunoprecipitated with anti-Flag antibody. Total lysates (input) and IP extracts were analyzed by Western blot with the indicated antibodies. **(B)**Flag-NSP2 IP in 4EHP-/- or GIGYF2-/- cells. Vectors encoding Flag-NSP2, or Flag as control, were transitory transfected in the Wild-Type (WT), 4EHP-/- or GIGYF2-/- HEK293 cells, and Flag IPs were performed in RNse A-treated extracts, followed by Western blot with the indicated antibodies. **(C)**Proximity Ligation Assay between Flag-NSP2 and endogenous GIGYF2. HEK293T cells were transfected with vector expressing Flag-NSP2, or Flag as control. PLA was performed using anti-FLAG and anti-GIGYF2 antibodies. Representative images of PLA (red), along with GIGYF2 immunofluorescence (IF; green) and DAPI (blue) are shown. Scale bars, 5 μm. **(D)**Representation of the NSP2 protein structure (PDB : 7MSW; (18)) and schematic cartoon of FLAG-tagged NSP2 truncations used in panel (E). **(E)**The N-terminal half of NSP2 binds 4EHP-GIGYF2. The indicated constructs were expressed in HEK293T and Flag IP were performed with RNase A-treated extracts. The starting material (Input) and bound fractions were analyzed by Western blot. EV: empty vector; FL: Full length.

To elucidate which protein is involved in the formation of this complex, GIGYF2- and 4EHP-Knock-Out (KO) HEK293 cells were used to conduct co-IPs with Flag-NSP2. Vectors expressing Flag-NSP2, or Flag as a control, were then transiently transfected in these KO populations, as well as in their Wild-Type (WT) counterpart. Following Flag IP in the 4EHP KO cells, we observed that the NSP2/GIGYF2 interaction was still detectable, while ZNF598 co-IP was reduced, indicating a plausible contribution of 4EHP in NSP2 binding. By contrast, in the absence of GIGYF2, we could not detect any interaction of NSP2 with either 4EHP or ZNF598 (Fig. 1B), supporting a central role of GIGYF2 in NSP2 binding. However, a contribution of 4EHP cannot be excluded since its level is decreased in the GIGYF2 KO cells due to a co-stabilization effect (Fig. 1B and (22)).

The sub-cellular localization of NSP2 was then examined using an *in situ* proximity ligation assay (PLA). PLA was conducted in Flag-NSP2-expressing cells and antibodies raised against the Flag sequence and endogenous GIGYF2 were used. Following confocal imaging, a spot-like signal was abundantly detected in the cytoplasm of HEK293T cells expressing Flag-NSP2, indicating the spatial proximity between endogenous GIGYF2 and Flag-NSP2 (Fig. 1C). This PLA signal was not observed in cells transfected with a control plasmid, supporting the specificity of this interaction. We then performed confocal live imaging to analyze the sub-cellular distribution of NSP2. Vectors expressing mCherry-tagged NSP2 fused either at the N- or C-terminal extremity were individually co-transfected along with an eGFP-tagged GIGYF2-encoding vector in HEK293T cells. A diffuse cytoplasmic signal was detected upon the expression of both eGFP-tagged GIGYF2 and mCherry-NSP2, indicating their cytoplasmic distribution (Supplementary Fig. 1A and B). The eGFP-GIGYF2 pattern was also punctuated by distinct condensates, in agreement with the presence of GIGYF2 in P-bodies, a subclass of RNA granules enriched in translationally repressed mRNAs and silencing factors (36). We did not find any co-localization between mCherry-fused NSP2 and the eGFP-GIGYF2 foci (Supplementary Fig. 1A and B), indicating that NSP2 could preferentially bind the diffuse form of GIGYF2 instead of its P-body-associated counterpart. To confirm this point, we took advantage of the properties of biotinylated-isoxazole (b-isox), a compound which selectively precipitates proteins located in RNA granules such as P-bodies and stress granules (37,38) (Supplementary Fig. 1C). Extracts of HEK293T expressing Flag-NSP2 were exposed to 100 μM of b-isox, or DMSO as a mock control. By comparing the level of proteins left in the soluble fraction after precipitation (unbound) with the precipitated fractions (pellet), we confirmed that Flag-NSP2 was not recovered in the pellet whereas a significant proportion of endogenous GIGYF2 was selectively enriched in the b-isox precipitate, alongside DDX6, a known resident of P-bodies (Supplementary Fig. 1D). Since NSP2 could not be precipitated by b-isox along with GIGYF2, we concluded that it only binds the diffuse form of GIGYF2 in the cytoplasm.

We next investigated which part of NSP2 binds the 4EHP-GIGYF2 complex. For this purpose, we generated a collection of vectors encoding Flag-tagged truncated versions of NSP2 which were expressed in HEK293T for Flag IP (Fig. 1D and E). Incremental terminal deletions revealed that the N-terminal half of NSP2 is required for the maximal co-IP of the endogenous 4EHP-GIGYF2 complex as well as ZNF598. While weakly expressed, the NSP2^1-350^ fragment remained the minimal segment which was able to bind 4EHP-GIGYF2. Interestingly, deleting either the N-terminal extremity (fragment NSP2^107-350^) or the middle region (fragment NSP2^1-212^) impaired the NPS2/4EHP-GIGYF2 interaction, indicating a large interaction surface involving an intact N-terminal half of NSP2 (Fig. 1E).

### The NSP2/GIGYF2-4EHP interaction involves multiple binding sites

We then sought to delineate which part of the 4EHP-GIGYF2 complex is targeted by NSP2. Human GIGYF2 is a 150 kDa scaffolding protein composed of several domains interspaced by intrinsically disordered regions. These include the 4EHP-binding domain at the N-terminus, the so-called GYF domain, a putative Single Alpha-Helix (SAH) and many glutamine-rich stretches (polyQ) at the C-terminus (Fig. 2A)(22,39–41). V5-tagged fragments of GIGYF2 were designed to isolate these features, and expressed along with Flag-NSP2 for Flag IP (Fig. 2A and B). Our WB analysis of the IP fractions revealed that NSP2 binds two distinct segments of GIGYF2, namely the central putative SAH region (residues: 742-1,085), and to a smaller extent, the N-terminal extremity containing the 4EHP-binding motif (residues: 1-267; Fig. 2B). The strong interaction between NSP2 and GIGYF2^742-1,085^ prompted us to test their direct association. Full-length hexahistidine (His6)-tagged NSP2 and Glutathione S-transferase (GST)-fused GIGYF2^742-1,085^ were recombinantly expressed in *Escherichia coli* (*E. coli*), and individually purified to perform a His pull-down assay. Following incubation of His_6_-NSP2 with untagged GIGYF2^742-1,085^ on a Ni-NTA resin, analysis of the eluates by SDS-PAGE and Coomassie blue staining revealed a specific retention of GIGYF2^742-1,085^ with His_6_-NSP2, confirming their direct interaction *in vitro* (Fig. 2C). GST alone was used as control and did not show any pull-down by His_6_-NSP2, supporting the specificity of the NSP2-GIGYF2 interaction. We also successfully detected this interaction when both His_6_-NSP2 and GST-GIGYF2^742-1,085^ were co-expressed in *E.coli*. In this case, a GST pull-down assay was performed using an *E. coli* lysate and a specific His_6_-NSP2 retention on GST-GIGYF2^742-1,085^-bound beads was detected by both Coomassie blue staining and WB (Supplementary Fig. 2). Similarly, we sought to test the capacity of His_6_-NSP2 to bind the N-terminal region of GIGYF2 (residues: 1-267), but failed to obtain this recombinant GIGYF2 fragment with a sufficient yield and solubility (data not shown).

**Figure 2.**
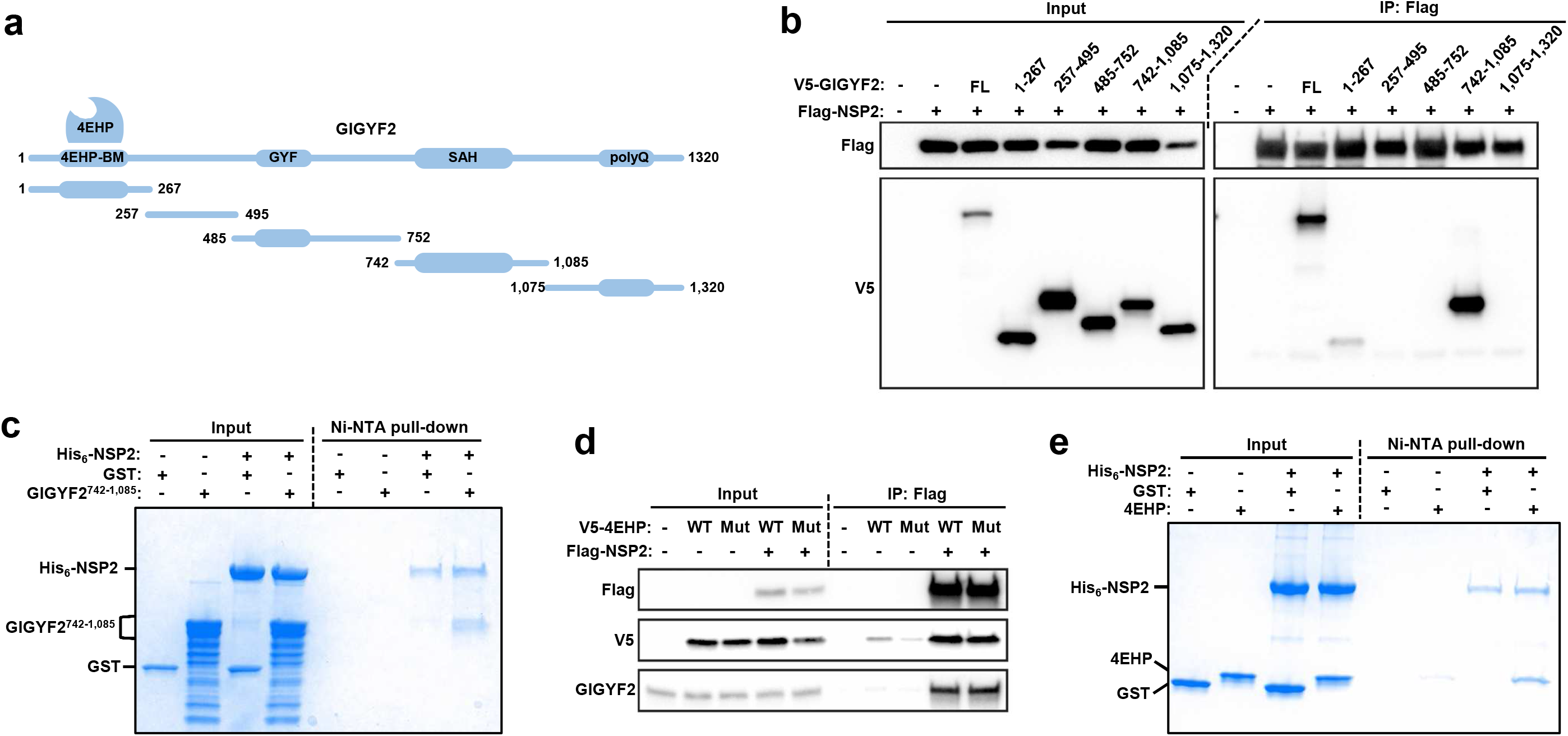
The NSP2/GIGYF2-4EHP interaction involves multiple binding sites. **(A)**Schematic cartoon of the V5-tagged GIGYF2 fragments used in panel (B). 4EHP-BM: 4EHP-binding motif; GYF: glycine-tyrosine-phenylalanine domain; SAH: putative single alpha helix; polyQ: glutamine-rich stretches. **(B)**Western blot showing the interaction between Flag-NSP2 and two regions in GIGYF2. Vectors expressing Flag-NSP2 and the indicated fragments of V5-tagged GIGYF2 were transitory transfected in HEK293T to perform Flag IP. Extracts were RNase A-treated. Empty vectors were used as negative controls (-). **(C)**Ni-NTA pull-down assay showing the interactions between recombinant His_6_-NSP2 and untagged GIGYF2^742-1,085^. GST served as negative control. The starting material (Input) and bound (NiNTA pull-down) fractions were analyzed by SDS-PAGE followed by Coomassie blue staining. **(D)**Western blot showing the interaction between Flag-NSP2 and V5-4EHP in a GIGYF2-independent manner. RNase A-treated extracts from cells expressing Flag-NSP2 along with V5-4EHP, WT or carrying the W95A substitution (Mut), were used for Flag IP. Inputs and bound fractions were analyzed by Western blotting using the indicated antibodies. Empty vectors served as negative controls (-). **(E)**Ni-NTA pull-down assay showing the interactions between recombinant His_6_-NSP2 and untagged full-length 4EHP. GST served as negative control and the indicated extracts were analyzed by SDS-PAGE followed by Coomassie blue staining.

Since an interaction was detected between NSP2 and the 4EHP-binding region of GIGYF2 (residues: 1-267) by co-IP, we speculated that 4EHP could also bind NSP2 independently of GIGYF2. To test this, we used a V5-tagged version of 4EHP carrying the W95A substitution which is known to disrupt its interaction with GIGYF2 (39). HEK293T cells were co-transfected by vectors encoding Flag-NSP2 and V5-4EHP^W95A^, or its WT counterpart. Using Flag IPs, we found that both WT and W95A versions of 4EHP were co-immunoprecipitated by NSP2 to a comparable extent (Fig. 2D), indicating that 4EHP can bind NSP2 independently of GIGYF2. This result prompted us to evaluate their direct association *in vitro*. As performed with GIGYF2^742-1,085^, full-length GST-fused 4EHP was expressed in *E. coli*, and the purified untagged protein was incubated with His_6_-NSP2 to perform a His pull-down assay. In agreement with our co-IP results, we found a specific binding of 4EHP to His_6_-NSP2, thus confirming their interaction in a GIGYF2-independent manner (Fig. 2E).

### NSP2 reduces the silencing capacity of GIGYF2

The role of 4EHP-GIGYF2 as a translational repressor has been described in various contexts (for review, see (21)). Having shown that NSP2 directly targets the 4EHP-GIGYF2 complex through at least two contact points, namely the region 742-1,085 of GIGYF2 and 4EHP itself, we wished to assess whether the silencing capacity of 4EHP-GIGYF2 could be altered by NSP2. To address this point, we used the λN-BoxB tethering approach in HEK293T cells (42). A *Renilla* luciferase reporter mRNA containing five BoxB sequences (RLuc-5BoxB) in the 3’ untranslated region (3’UTR) was expressed with a plasmid expressing V5-tagged GIGYF2 fused to a λN peptide, which has a high affinity for the BoxB sequences (Fig. 3A). The silencing capacity of GIGYF2 was assessed following the measurement of luciferase activity in HEK293T expressing Flag-NSP2 or Flag as control. The expression of Flag-NSP2 and λN-GIGYF2 was also verified by WB (Fig. 3C). As expected, we observed that the recruitment of GIGYF2 to the 3’ UTR markedly reduced luciferase activity in control cells (5-fold repression). Interestingly, this λN-GIGYF2-mediated repression was reduced upon expression of Flag-NSP2 (3-fold repression), indicating an inhibitory effect of NSP2 on GIGYF2 function in silencing (Fig. 3A).

**Figure 3.**
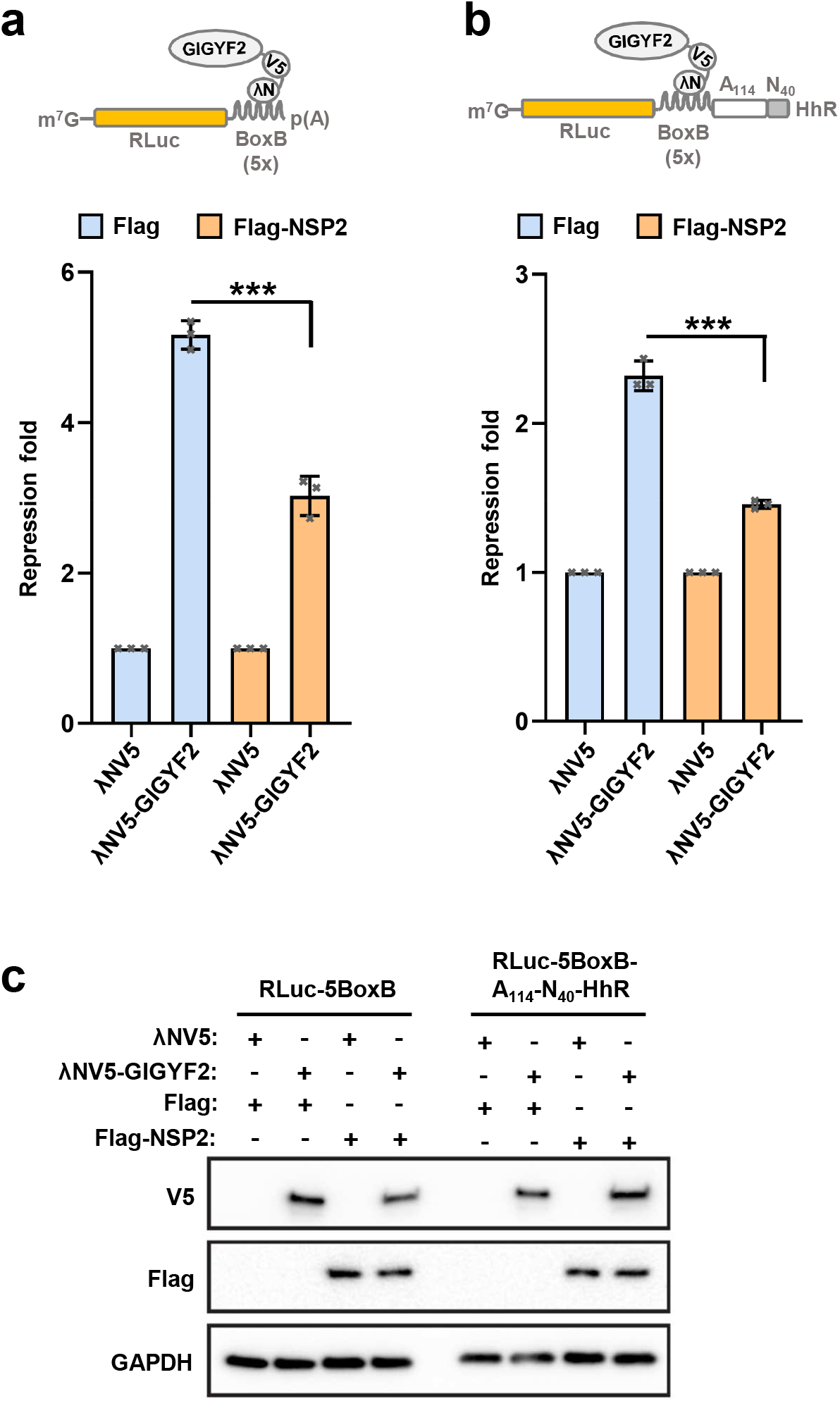
NSP2 decreases the silencing capacities of GIGYF2 *in cellulo*. **(A)**Artificial tethering of GIGYF2 to the 3’ UTR of a reporter mRNA. The upper panel shows a schematic of the λN/BoxB tethering assay with the RLuc-5boxB reporter construct. Recruitment of GIGYF2 to the Renilla Luciferase (RLuc) mRNA was mediated by the fused λN peptide. Rluc luminescence was normalized against Firefly luciferase (FLuc) level, and repression fold was calculated by dividing the relative luciferase activity of the cells transfected with the control pCI-λNV5 vector (λNV5) by the luciferase activity of λN-GIGYF2 expressing cells. The mean values (±SD) from three independent experiments are shown and the *P* value was determined by two-tailed Student’s t-test: (***) P < 0.001. **(B)**Artificial tethering of GIGYF2 to the 3’ UTR of a reporter mRNA which is refractory to deadenylation. The upper panel shows a schematic of the RLuc-5boxB-A_114_-N_40_-HhR reporter. HEK293T cells were co-transfected with vectors expressing either λNV5-GIGYF2, or λNV5 as a control, along with RLuc-5boxB-A_114_-N_40_-HhR and FLuc. Vectors encoding Flag-NSP2 or Flag (empty vector) were also added in the transfection mixture. RLuc luminescence was normalized against the FLuc level and analyzed as in (A). **(C)**Extracts from the HEK293T cells used in (A) and (B) were analyzed by Western blot with the indicated antibodies. GAPDH was used as a loading control.

Several studies have reported that GIGYF2 has two distinct mechanisms of repression: one is 4EHP-dependent and affects translation; the other is 4EHP-independent and involves the deadenylase activity of the CCR4-NOT complex (36). Our co-IP data showed that NSP2 did not interact with CCR4-NOT (Fig. 1A), suggesting that it should preferentially impair the 4EHP-dependent activity of GIGYF2. To test this, λN-GIGYF2 was tethered to a RLuc-5BoxB mRNA containing a self-cleaving hammerhead ribozyme (HhR) at the 3’-end to generate a poly(A) stretch of 114 nucleotides, followed by 40 nucleotides to block CCR4-NOT-dependent deadenylation (RLuc-5BoxB-A_114_-N_40_-HhR), as previously described (18). When tethered to this reporter, GIGYF2 induced a 2.2-fold repression of this reporter in cells transfected with a control vector (Fig. 3B). By contrast, we observed that NSP2 expression decreased the GIGYF2-mediated repression of the reporter to 1.4-fold, indicating a specific role of NSP2 in blocking the deadenylation-independent role of GIGYF2.

### NSP2 impairs miRNA-mediated silencing

The miRNA-induced translational repression is effected by the 4EHP-GIGYF2 complex in human cells (27,28,30). To evaluate the impact of NSP2 in miRNA-mediated silencing, we transiently transfected HEK293T cells with a *Renilla* luciferase construct either lacking (RLuc), or containing six bulged *let7a* miRNA binding sites in its 3’ UTR (RLuc-*6let7a*), together with a firefly luciferase construct (FLuc) as a transfection control (Fig. 4A). Normalized RLuc activity was markedly reduced by the presence of *let7a*-binding sites, with ~40-fold repression in cells expressing a control vector. By contrast, expression of NSP2 resulted in a decreased repression magnitude of the RLuc-*6let7a* reporter compared to RLuc (~30-fold), indicating a reduced *let7a*-mediated silencing in the presence of NSP2 (Fig. 4A).

**Figure 4.**
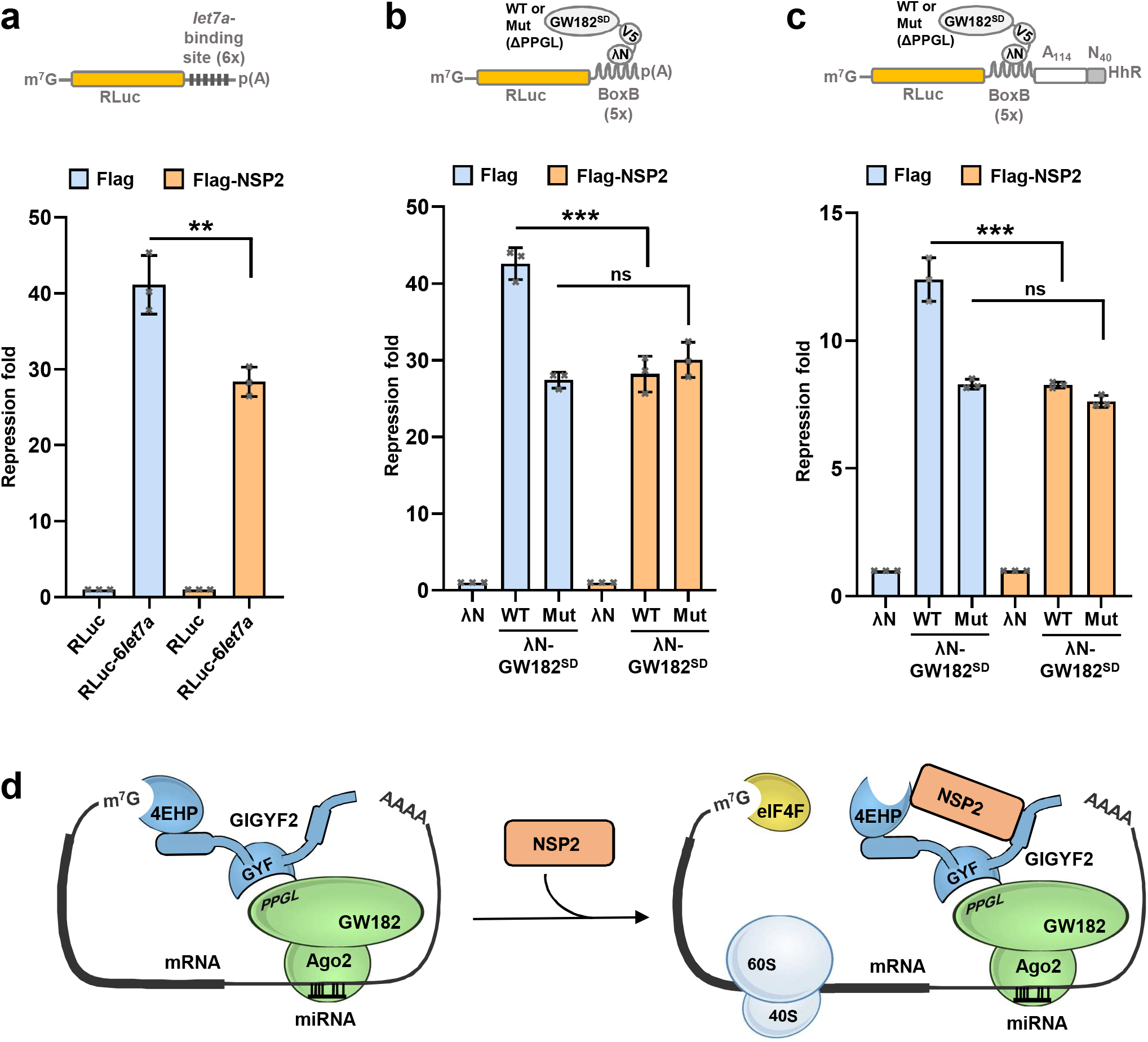
NSP2 impairs miRNA-mediated silencing. **(A)**NSP2 decreases *let7a*-mediated silencing. The upper panel shows a schematic of the RLuc-*6let7a* reporter. HEK293T cells were co-transfected with RLuc or RLuc-*6let7a* plasmids, along with FLuc construct to account for variations in transfection efficiency. Vectors encoding Flag-NSP2 or Flag (empty vector) were also added in the transfection mixture. Repression fold was calculated by dividing the relative luciferase activity of the cells transfected with the RLuc vector by the luciferase activity of RLuc-*6let7a* expressing cells. Error bars indicate ±SD (*n* = 3). (**) *P* < 0.01 (two-tailed Student’s t test). **(B)**Artificial tethering of GW182^SD^ to the 3’ UTR of a reporter mRNA. The upper panel shows a schematic of the λN/BoxB tethering assay with the RLuc-5boxB reporter construct. HEK293T cells were co-transfected with vectors expressing either λNV5-GW182^SD^, WT or a ΔPPGL mutant (Mut), or λNV5 as a control, along with RLuc-5boxB and FLuc. Vectors encoding Flag-NSP2 or Flag (empty vector) were also added in the transfection mixture. RLuc luminescence was normalized against the FLuc level. The mean values (±SD) from three independent experiments are shown and the *P* value was determined by two-tailed Student’s t-test: (ns) non-significant, (***) *P* < 0.001. **(C)**Artificial tethering of GW182^SD^ to the 3’ UTR of a reporter mRNA which is refractory to deadenylation. The upper panel shows a schematic of the RLuc-5boxB-A_114_-N_40_-HhR reporter. Transfections were performed as in (B), except that GW182^SD^ was tethered on the RL-5*boxB*-A_114_-N_40_-HhR reporter. Data are presented as mean ± SD (n = 3). (***) *P* < 0.001 (two-tailed Student’s t test). **(D)**Model of NSP2-mediated negative regulation of miRNA function. In absence of NSP2 (left panel), miRISC recruits the 4EHP-GIGYF2 complex to effect translational silencing of the targeted mRNA. Following NSP2 expression (right panel), the function of 4EHP-GIGYF2 is physically targeted by NSP2 and its silencing capacity is impaired. The assembly of this complex subsequently alters the magnitude of miRNA-induced silencing.

Evidence suggests that the 4EHP-GIGYF2 complex contributes to miRNA action through the binding of GIGYF2 to a proline-rich motif (PPGL) in the silencing domain (SD) of GW182, the scaffolding protein of miRISC (30). To investigate whether NSP2 could impair GW182-mediated silencing, the silencing domain of GW182 was artificially tethered to the RLuc-5BoxB reporter in control and Flag-NSP2-expressing cells. We found that tethering GW182^SD^ induced a ~40-fold repression of the RLuc-5BoxB level in the control cells, while this silencing effect was reduced in NSP2-expressing cells (~30-fold; Fig. 4B). As a control, a GW182^SD^ mutant carrying a deletion of the GIGYF2-binding motif (ΔPPGL) was used to evaluate the contribution of GIGYF2. Tethering this ΔPPGL mutant on RLuc-5BoxB in control and Flag-NSP2 cells showed that its silencing activity is not affected by NSP2 expression, indicating that NSP2 action on GW182^SD^-mediated silencing is exerted through targeting GIGYF2 (Fig. 4B). We detected a comparable result when GW182^SD^ was tethered on the RLuc-5BoxB-A_114_-N_40_-HhR reporter (Fig. 4C). Similarly, the silencing activity of its ΔPPGL version was unchanged upon NSP2 expression, confirming a specific alteration of the deadenylation-independent role of GIGYF2 by NSP2. Altogether, these results indicate that the GIGYF2-dependent repressive activity of miRISC is altered upon expression of NSP2 in human cells.

## DISCUSSION

Upon infection, SARS-CoV-2 impairs splicing, export, translation and degradation of host mRNAs (43,44). Here, we present evidence to support a new layer of complexity in the post-transcriptional alteration of the host transcriptome by SARS-CoV-2. We propose that SARS-CoV-2 NSP2 directly targets the 4EHP-GIGYF2 complex to impair the silencing capacity of miRNAs (Fig. 4D). This mechanism therefore unveils the potential impact of NSP2 on the post-transcriptional silencing of gene expression of human cells, pointing out 4EHP-GIGYF2 targeting as a possible strategy of SARS-CoV-2 to take over the silencing machinery and to suppress host defenses.

How does NSP2 impair 4EHP-GIGYF2 function? Combining co-IP experiments and *in vitro* binding assays with recombinant proteins, we concluded that NSP2 uses its N-terminal region encompassing its conserved zinc finger domain, to interact with the 4EHP-GIGYF2 complex. Our pull-down assays indicate the direct interaction of NSP2 with both 4EHP and two domains from GIGYF2, confirming a sophisticated mode of binding *in cellulo*. While we searched for the minimal region of NSP2 required for these interactions, we failed to narrow down a fragment smaller than the 1-350 region since truncations at both extremities of this domain abrogate its binding to 4EHP-GIGYF2 (Fig. 1E). Nevertheless, this remains in agreement with Gupta *et al.* who pointed out that the G262V and G265V mutations located within this region of NSP2 disrupted binding to 4EHP-GIGYF2 (18). It now remains to be determined whether NSP2 binding induces conformational changes in 4EHP-GIGYF2 that impair either the cap-binding pocket of 4EHP, or influence the recruitment of GIGYF2 co-factors such as CCR4-NOT and DDX6. Further investigation will be needed to fully resolve the structural basis of the NSP2/4EHP-GIGYF2 complex and thus elucidate NSP2 action on miRNA-mediated silencing.

At the moment it is uncertain what the functional interplay between SARS-CoV-2 infection and miRNA-mediated silencing is in human cells. Host miRNAs are known to be produced as a part of antiviral response to counteract the infection by targeting viral transcripts, although SARS-CoV-2 infection was recently shown to have minimal impact on the miRNA repertoire of its host cell (45–47). Computational analyses have predicted the presence of many putative miRNA-binding sites on the SARS-CoV-2 genome, suggesting that the SARS-CoV-2 genome could be actively targeted by host miRNAs (48–50). Particularly worth mentioning is the work of Xie *et al.* which recently identified *let-7* binding sites in the coding sequence of S and M proteins of SARS-CoV-2 genome, and experimentally confirmed that *let-7* blocks SARS-CoV-2 replication by targeting S and M proteins (48). Through the NSP2/4EHP-GIGYF2 axis, SARS-CoV-2 could therefore escape from the host defense system by impairing the function of the effector machinery of miRNAs. The recent discovery that 4EHP and GIGYF2 are needed for infection by SARS-CoV-2 could reinforce this idea, although further research will be required to test this possibility (8).

The silencing capacity of miRISC is partially impeded upon NSP2 expression, as demonstrated by the de-repression of GW182-or *let-7a*-mediated inhibition of a reporter mRNA in NSP2-expressing cells (Fig. 4A-C). This alteration can be extrapolated to other miRNAs whose action relies on 4EHP, such as *miR-145* or *miR-34a* (31,32). Recent evidence demonstrated that the 4EHP/*miR-34a* axis is required for the translational repression of mRNAs encoding IFN-β through targeting the 3’UTR of *Ifnb1* mRNA (31). Beyond miRNA, 4EHP-GIGYF2 also controls the production of TTP-targeted mRNAs that encode inflammatory cytokines such as TNF-α and IL-8 (22–26). In this context, a possible consequence of NSP2 expression could be the overproduction of early response pro-inflammatory cytokines. Exploring this point would be of utmost importance since impaired type I interferon activity and inflammatory responses are detected in severe COVID-19 patients (51). To examine whether NSP2 could impact the function of 4EHP in regulating IFN-β expression, we expressed NSP2 along with a reporter construct containing the 3’ UTR of *Ifnb1* mRNA into HEK293T cells (Supplementary Fig. 3A). Remarkably, the reporter expression was repressed ~2.9-fold in control cells, but only ~1.6-fold in NSP2-expressing cells, indicating that NSP2 could potentially unbalance the production of IFN-β through the *Ifnb1* 3’ UTR (Supplementary Fig. 3B). With this in mind, further investigations into whether the NSP2/4EHP-GIGYF2 axis can dysregulate sustained cytokine production may therefore prove useful.

In conclusion, our study raises the possibility that SARS-CoV-2 could target the human 4EHP-GIGYF2 complex to selectively modulate the expression of host and/or viral transcripts. These results open new directions to investigate the mechanisms underlying the pathogenicity of SARS-CoV-2 and how NSP2 could become a target for therapeutic intervention. Thus, our biochemical approaches and reporter-based assays may represent a novel framework to imagine innovative therapeutic approaches based on the interaction of NSP2 with 4EHP-GIGYF2.

## Materials and methods

### Cell culture

HEK293T cells (Sigma-Aldrich) were routinely maintained in high glucose Dulbecco’s Modified Eagle’s Medium (DMEM) with GlutaMAX supplemented with 10 % fetal bovine serum (FBS) and 2 % penicillin/streptomycin in a humidified atmosphere of 5 % CO2 at 37 °C. Flp-In T-REx 293 cells (Thermo Fisher Scientific) were grown in similar conditions supplemented with 100μg/ml zeocin and 15μg/ml blasticidin. The absence of mycoplasma contamination in cells was routinely tested. The cell line inducibly expressing Flag-NSP2 was generated by co-transfecting pcDNA5-FRT-TO-FH-Nsp2 (Addgene plasmid 157683) and pOG44 (Thermo Fisher Scientific) with a 1:10 ratio. Transfected cells were selected and maintained in media supplemented with 100μg/ml hygromycin. Expression of tagged proteins was induced for 24 h by addition of tetracycline to 1 μg/mL final concentration.

### Plasmid cloning

For Flag-NSP2 IP, pcDNA5-FRT-TO-FH-Nsp2 (Addgene plasmid 157683) was used. Truncated versions of NSP2 were generated by replacing the sequence encoding full-length NSP2 by PCR-amplified fragments at the BamHI-XhoI sites of the pcDNA5-FRT-TO-FH vector. The mCherry2-NSP2 expressing plasmids were created by cloning a PCR-amplified NSP2 fragment into the mCherry2-C1 (Addgene plasmid 54563) and mCherry2-N1 vectors (Addgene plasmid 54517) using EcoRI and BamHI restriction enzymes. To generate the pCI-λNV5-GIGYF2 vector, a fragment containing the GIGYF2 sequence was obtained by PCR from the pcDNA4/TO/GFP-GIGYF2 vector (Addgene plasmid 141189) (34) and inserted into the pCI-λNV5 at the XhoI-NotI sites. A similar strategy was used to generate the V5-tagged fragments of GIGYF2 using primers to isolate the following domains encompassing residues: 1-267; 257-495; 485-752; 742-1,085; 1,075-1,320. pCI-eGFP-GIGYF2 was generated by inserting the eGFP sequence, PCR-amplified from pEGFP-C1 (Clontech), at the NheI-XhoI sites of the pCI-Neo vector and then by adding the fragment encoding GIGYF2 at the XhoI-NotI sites. The full-length cDNA encompassing the coding region of human 4EHP, obtained by RT-PCR using total RNA from HEK293T cells, were cloned into the pCI-Neo vector (Promega) at the XhoI-NotI sites in frame with a sequence encoding a V5 tag inserted at the NheI-XhoI sites. To generate the pCI-λNV5-GW182^SD^ vector, a fragment encoding the residues 1382-1690 was cloned by PCR as a XhoI-NotI fragment from the pFRT/TO/FLAG/HA-DEST TNRC6C vector (Addgene plasmid 19885) (52) and inserted into the pCI-λNV5 at the XhoI-NotI sites. For recombinant GST-fused protein expression, full-length 4EHP and GIGYF2^742-1,085^ coding sequences contained in PCR-amplified BamHI-NotI and XhoI-NotI fragments, respectively, were cloned into a pGEX-6P-1 vector (Amersham) in frame with the GST coding sequence. The pET-28b vector (EMD Biosciences) was used to express NSP2 as a recombinant protein fused to an His_6_ tag at the N-terminus. For this, a PCR-amplified fragment encoding NSP2 was inserted at the XhoI-BamHI sites of pET-28b. Amino acid substitutions and deletions were introduced by site-directed mutagenesis using the QuikChange Kit (Agilent).

### Extract Preparation and Immunoprecipitation

Cells were resuspended in a lysis buffer containing 20 mM HEPES-KOH, pH 7.4, 100 mM NaCl, 2.5 mM MgCl2, 0.5 % NP40, 0.25 % sodium deoxycholate, supplemented with cOmplete EDTA-free Protease inhibitor and phosphatase inhibitor Cocktails (Roche), and incubated for 20 min on ice. The lysate was clarified by centrifugation at 10,000 × g for 10 min at 4 °C. One milligram of extract was used for immunoprecipitation with the indicated antibodies. Thirty microliters of pre-equilibrated Anti-FLAG L5 Magnetic Beads (Thermo Fisher Scientific, A36798) and RNase A (Thermo Fisher Scientific) were added, and the mixtures were rotated overnight at 4 °C. Beads were washed five times with lysis buffer and directly resuspended in protein sample buffer for Western blot analysis.

### Western blot and antibodies

Proteins were separated by SDS-PAGE on 4 – 15 % gradient gels (BioRad) and transferred onto PVDF membranes. The membranes were blocked in PBS containing 5 % non-fat milk and 0.1 % Tween 20 for 30 min at room temperature. Blots were probed with the following antibodies: rabbit anti-eIF4E2/4EHP (Proteintech, 12227-1-AP), rabbit anti-DDX6 (Proteintech, 14632-1-AP), rabbit anti-CNOT9 (Proteintech, 22503-1-AP), mouse anti-Flag M2 (Sigma-Aldrich, F1804), rabbit anti-GIGYF2 (Proteintech, 24790-1-AP), mouse anti-GAPDH (Proteintech, 60004-1-Ig), mouse anti-V5 tag (Invitrogen, R96025), rabbit anti-ZNF598 (Thermo Fischer Scientific, 703601) and mouse anti-6xHis Tag (Thermo Fischer Scientific, MA1-21315-HRP).

### Biotinylated isoxazole (b-isox)-mediated precipitation

HEK293T cells were resuspended in a lysis buffer (50 mM HEPES pH 7.5, 150 mM NaCl, 0.1% NP-40, 1 mM EDTA, 2.5 mM EGTA, 10% glycerol, 1 μM DTT) supplemented with cOmplete EDTA-free Protease inhibitor, and phosphatase inhibitors, and incubated for 20 min on ice. The lysate was clarified by centrifugation at 10,000 × g for 10 min at 4 °C. 50μg of the sample were mixed with 100 μM of b-isox (Sigma-Aldrich) and rotated at 4°C for 90 min. The incubated reaction was then centrifuged at 10,000 × g for 10 min to pellet the precipitates. The pellet was washed twice in the lysis buffer and resuspended in protein sample buffer for Western blot analysis. Proteins in the supernatant fractions were precipitated by addition of four volumes of cold acetone, incubated for 1 h at −20°C and centrifuged at 15,000 × g for 10 min to pellet the precipitates.

### CRISPR/cas9-mediated genome editing

Generation of 4EHP KO cells were described in Jafarnejad *et al.* (32). CRISPR-Cas9-mediated genome editing of HEK293 cells was performed according to Ran *et al.* (53). The following DNA oligonucleotides encoding a small guide RNA (sgRNA) cognate to the coding region of *Gigyf2* gene were used: 5’-CACCGGGAGGAACCCCTTCCACCAT and 5’-AAACATGGTGGAAGGGGTTCCTCCC. These oligos which contain BbsI restriction sites were annealed creating overhangs for cloning of the guide sequence oligos into pSpCas9(BB)-2A-Puro (PX459) V2.0 (Addgene plasmid 62988) by BbsI digestion. To generate KO HEK293 cells, we transfected 700,000 cells with the pSpCas9(BB)-2A-Puro plasmid. 24 hr after transfection, puromycin was added in the cell medium. After 72 h, puromycin-resistant cells isolated into 96-well plates and cultivated until monoclonal colonies were obtained. Clonal cell populations were analyzed by WB for protein depletion.

### PLA and confocal microscopy scanning

PLA was performed using the Duolink kit (Sigma-Aldrich) according to manufacturer’s instructions. Transfected cells (100,000) were fixed in 4 % paraformaldehyde for 20 min, washed twice in PBS, permeabilized in PBS containing 0.1% Triton X100 for 5 min and incubated with primary antibodies for 1 h at 37°C in a pre-heated humidity chamber. After two washes in PBS-Tween 0.1%, cells were incubated with the appropriate PLA probes for 1 h at 37°C then washed in PBS-T, and the ligation solution was added on the coverslips and incubated for 30 min at 37°C. Finally, the amplification solution containing a DNA polymerase was added and incubated with the cells for 100 min at 37°C. For simultaneous immunofluorescence labelling of GIGYF2, secondary antibody coupled with Alexa Fluor 488 (Thermo Fisher Scientific) was added in the amplification solution. After final washes, the cells were mounted on glass slides in a mounting solution with DAPI, and imaging was performed on a LEICA-SP8ST-WS confocal microscope. For live confocal staining, HEK293T cells were grown in U-Slide ibiTreat coverslip multi-well chambers and transfected with 20 ng of vectors encoding the indicated proteins. Fluorescence was visualized under a LEICA-SP8ST-WS confocal microscope using resonant scanning.

### Expression and purification of His_6_-NSP2

His_6_-NSP2 was expressed in *E. coli* BL21 (DE3) Gold (Agilent technologies). Large-scale expression was done in 1 L of auto-inducible terrific broth media (ForMedium AIMTB0260) supplemented with kanamycin (50 μg/ml), first at 37°C for 3 h and then at 18 °C overnight. The cells were harvested by centrifugation at 4,000 rpm for 30 min and the pellets were resuspended in 30 ml of lysis buffer (50 mM HEPES pH 7.5, 300 mM NaCl, 10 μM ZnCl_2_, 2 mM MgCl_2_ and 20 mM Imidazole) supplemented with 1 protease inhibitor tablet (Roche), 0.5 mM PMSF and 30 μL benzonase nuclease (Millipore Sigma). The cells were lysed by sonication on ice and the lysate clearance was performed by centrifugation at 20,000 g for 30 min. The supernatant was applied on Ni-NTA resin pre-equilibrated with the lysis buffer, incubated at 4 °C on a rotating wheel for 1 h, followed by a washing step with 30 mL of washing buffer (50 mM HEPES pH 7.5, 1 M NaCl and 10 μM ZnCl_2_). His_6_-NSP2 was eluted by the addition of 15 ml of elution buffer (50 mM Tris/HCl pH 8.0, 300 mM NaCl, 10 μM ZnCl_2_, 2 mM MgCl_2_ and 300 mM Imidazole), followed by concentrating up to 10 ml by a 30 kDa cutoff concentrator. The sample was then diluted to 50 mL using Heparin buffer A (50 mM Tris/HCl pH 8.0, 5 mM β-mercaptoethanol, 10 μM ZnCl_2_), injected on a 5 mL Heparin HP column (Cytiva) and eluted using a NaCl linear gradient from 75 mM (7.5 % Heparin buffer B: 50 mM Tris/HCl pH 8.0, 1 M NaCl, 5 mM β-mercaptoethanol, 10 μM ZnCl_2_) to 1 M (100 % Heparin buffer B). The fractions containing His_6_-NSP2 protein were collected and concentrated up to 5 ml, followed by injection on a Superdex 200 increase 10/300 size-exclusion column (Cytiva) with Gel filtration buffer (50 mM HEPES pH 7.5, 250 mM NaCl, 5 mM β-mercaptoethanol and 10 μM ZnCl_2_). The fractions containing His_6_-NSP2 were collected and concentrated.

### Expression and purification of 4EHP and GIGYF2^742-1,085^

Expression of GST-4EHP and GST-GIGYF2^742-1,085^ was carried out in *E. coli* 3 BL21 (DE3) Codon+ (Novagen) in 1 L of auto-inducible terrific broth media (ForMedium AIMTB0260) supplemented with ampicillin at 100 μg/mL and chloramphenicol at 25 μg/mL. When the OD_600 nm_ reached 0.6–0.8, cultures were incubated 20 °C for 20 h. Bacteria were harvested by centrifugation and resuspended in lysis buffer (20 mM HEPES pH 7.5, 200 mM NaCl, 5 mM β-mercaptoethanol). Cell lysis was performed by sonication on ice. After centrifugation for 30 min at 20,000 g, 4°C clarified samples were transferred to batch-bind with Glutathione SepharoseTM 4B (Cytiva) resin for ~1 hr at 4°C followed by a washing step with 30 mL of washing buffer (20 mM HEPES pH 7.5, 1 M NaCl and 5 mM β-mercaptoethanol) supplemented with 10 mM ATP, followed by an overnight 3C protease digestion in order to remove GST. The untagged GIGYF2^742-1,085^ domain was present in the flowthrough and further purified on an HiTrap S FF column (Cytiva) using a linear gradient of 92.5% Hitrap buffer A (20 mM HEPES pH7.5, 5 mM β-mercaptoethanol) to 100% Hitrap buffer B (20 mM HEPES pH 7.5, 1 M NaCl and 5 mM β-mercaptoethanol). The peak fractions corresponding to 4EHP and GIGYF2^742-1,085^ were pooled, concentrated and used for pull-down assays.

### Pull-down assays

Pull-down experiments were performed by incubating 1 nmol of His_6_-NSP2 with equimolar amount of untagged GIGYF2^742-1,085^ or 4EHP, or GST used as control. All proteins were free of nucleic acids according to the OD_280 nm_/OD_260 nm_ ratio. Binding buffer (20 mM HEPES pH 7.5, 200 mM NaCl, 50 mM imidazole, 10% glycerol and 0.1 % triton X-100) was added to a final volume of 60 μL. The reaction mixtures were incubated on ice for 1 h. 10 μL was withdrawn and used as an input fraction for SDS–PAGE analysis. The remaining 50 μL were incubated at 4 °C for 2 h with 40 μg of HisPur Ni–NTA magnetic beads (Thermo Scientific) pre-equilibrated in binding buffer, in a final volume of 200 μL. After binding, beads were washed three times with 500 μL of binding buffer. Bound proteins were eluted with 50 μl of elution buffer (20 mM HEPES pH 7.5, 200 mM NaCl, 250 mM imidazole, 10 % glycerol and 0.1 % triton X-100). Samples were resolved on SDS– PAGE and visualized by Coomassie blue staining.

### Co-expression and GST pull-down between His_6_-NSP2 and GST-GIGYF2^742-1,085^

Full-length His6-NSP2 and GST-GIGYF2^742-1,085^ proteins were co-expressed in BL21 (DE3) Gold *E. coli* (Agilent technologies). Small-scale expression was done in 5 ml of auto-inducible terrific broth media (ForMedium, AIMTB0260) supplemented with ampicillin (100 μg/ml) and kanamycin (50 μg/mL), first at 37°C for 3 h and then at 18°C overnight. The cells were harvested by centrifugation at 4,000 rpm for 30 min and the pellets were resuspended in 750 μl of lysis buffer (50 mM HEPES pH 7.5, 200mM NaCl, 5 mM β-mercaptoethanol). The cells were lysed by sonication on ice and the lysate clearance was performed by centrifugation at 13,000 g for 30 min. The supernatant was applied on Glutathione SepharoseTM 4B resin (Cytiva) pre-equilibrated with the lysis buffer, incubated at 4°C on a rotating wheel for 1 h, followed by a washing step with 1 ml of lysis buffer (50 mM HEPES pH 7.5, 200mM NaCl, 5 mM β-mercaptoethanol). Retained proteins were eluted by 50 μl of elution buffer (50 mM HEPES pH 7.5, 200mM NaCl, 5 mM β-mercaptoethanol, 20mM GSH), followed by SDS-PAGE and Coomassie blue staining or Western blot.

### Tethering and luciferase assays

Tethering assays were performed as previously described (27,54). Briefly, HEK293T cells were transfected with 20 ng of RLuc-5BoxB (or RLuc-5BoxB-A_114_-N_40_-HhR), 5 ng of FLuc, and 100 ng of λN-fusion constructs per well in a 24-well plate by using Lipofectamine 2000 (Thermo Scientific, 11668019) according to the manufacturer’s instructions. 100 ng of vectors encoding Flag-NSP2 or Flag (empty vector) were also added in the transfection mixture. Cells were lysed 24 h after transfection and luciferase activities were measured with the Dual-Luciferase Reporter Assay System (Promega) in a GloMax 20/20 luminometer (Promega). RL activity was normalized to the activity of co-expressed FL, and the normalized RL values are shown as repression fold relative to the indicated control. For experiments with miRNA reporters, HEK293T were co-transfected in a 24-well plate with 100 ng of vectors encoding Flag-NSP2, or Flag (empty vector) as control, 20 ng of RLuc-*6let7a* and 5 ng of FLuc plasmid. For the reporter containing the 3’UTR of *Ifnb1*, 20 ng of psiCHECK2-RLuc-*Ifnb1* 3’ UTR reporter (31), or empty psiCHECK2 (Promega), was added along with 100 ng of vectors encoding Flag-NSP2 or Flag (empty vector).

## Acknowledgments

We thank N. Ulrich for helpful technical assistance, and A. Garcia (Rockefeller University, New York) for sharing information on sgRNA sequences targeting the *Gigyf2* gene. W. Filipowicz (FMI, Basel) and N. Gehring (University of Cologne) are acknowledged for their gifts of pCI-RL (5BoxB or 6let7a) and pCI-λN-V5, respectively. pFRT/TO/FLAG/HA-DEST TNRC6C was a gift from Thomas Tuschl (Addgene plasmid 19885), pcDNA5-FRT-TO-FH-Nsp2 from David Tollervey (Addgene plasmid 157683) and pcDNA4/TO/GFP-GIGYF2 from Simon Bekker-Jensen (Addgene plasmid 141189). For mCherry2 fusion, mCherry2-C1 (Addgene plasmid 54563) and mCherry2-N1 (Addgene plasmid 54517) were gifts from Michael Davidson. The pSpCas9(BB)-2A-Puro vector was a gift from Feng Zhang (Addgene plasmid 62988). C.C. acknowledges financial supports from the Centre National pour la Recherche Scientifique (CNRS), Ecole Polytechnique, Les Entreprises contre le cancer – Gefluc – Paris Île-de-France, and Fondation ARC. L.Z. is supported by a PhD fellowship from the China Scholarship Council (CSC).

## Contributions

L.Z., C.M. and C.C. performed the experiments. L.Z., M.G. and C.C. designed research and analyzed the experiments, and L.Z., C.M., M.G. and C.C. wrote the paper.

---

**Supplementary Figure 1.**
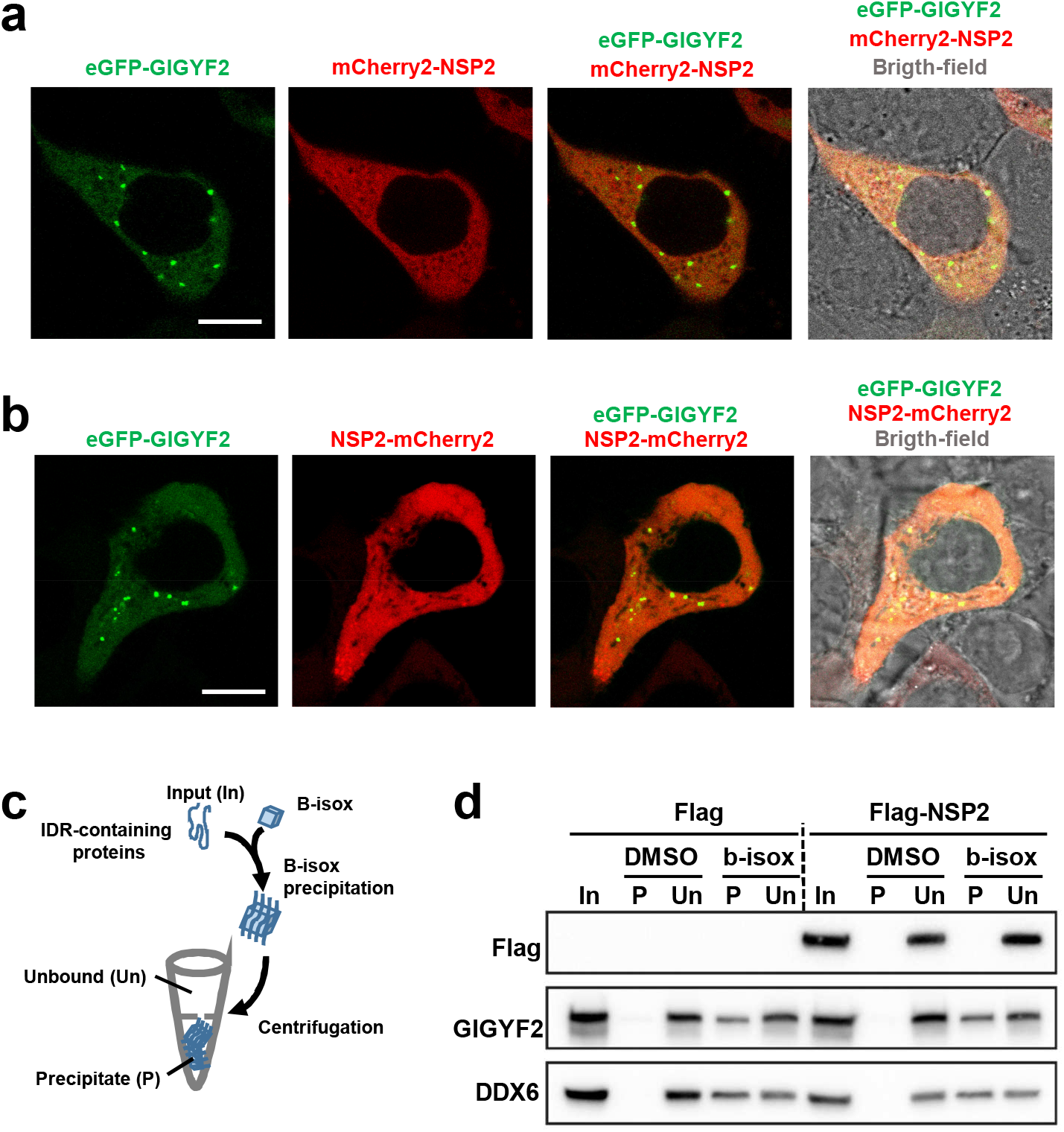
**(A-B)** Live cell confocal imaging of HEK293T cells transfected with plasmids expressing eGFP-GIGYF2 and NSP2 fused to mCherry2 at the N-terminus (A) or C-terminus (B). Z-projection of 3 stacks (0.4μm each); error bar = 10μm. **(C)**Schematic overview of the biotinylated isoxazole (b-isox)-mediated precipitation of Intrinsically-Disordered Region (IDR)-containing proteins. **(D)**B-isox precipitation of endogenous GIGYF2 in Flag-NSP2 expressing HEK293T cell extract. Cells transfected by an empty vector (Flag) were used as a negative control. Input (In), Precipitated (P) and Unbound (Un) fractions were analyzed by Western blot with the indicated antibodies. DMSO, used as the solubilizing agent for b-isox, served as a mock control.

**Supplementary Figure 2.**
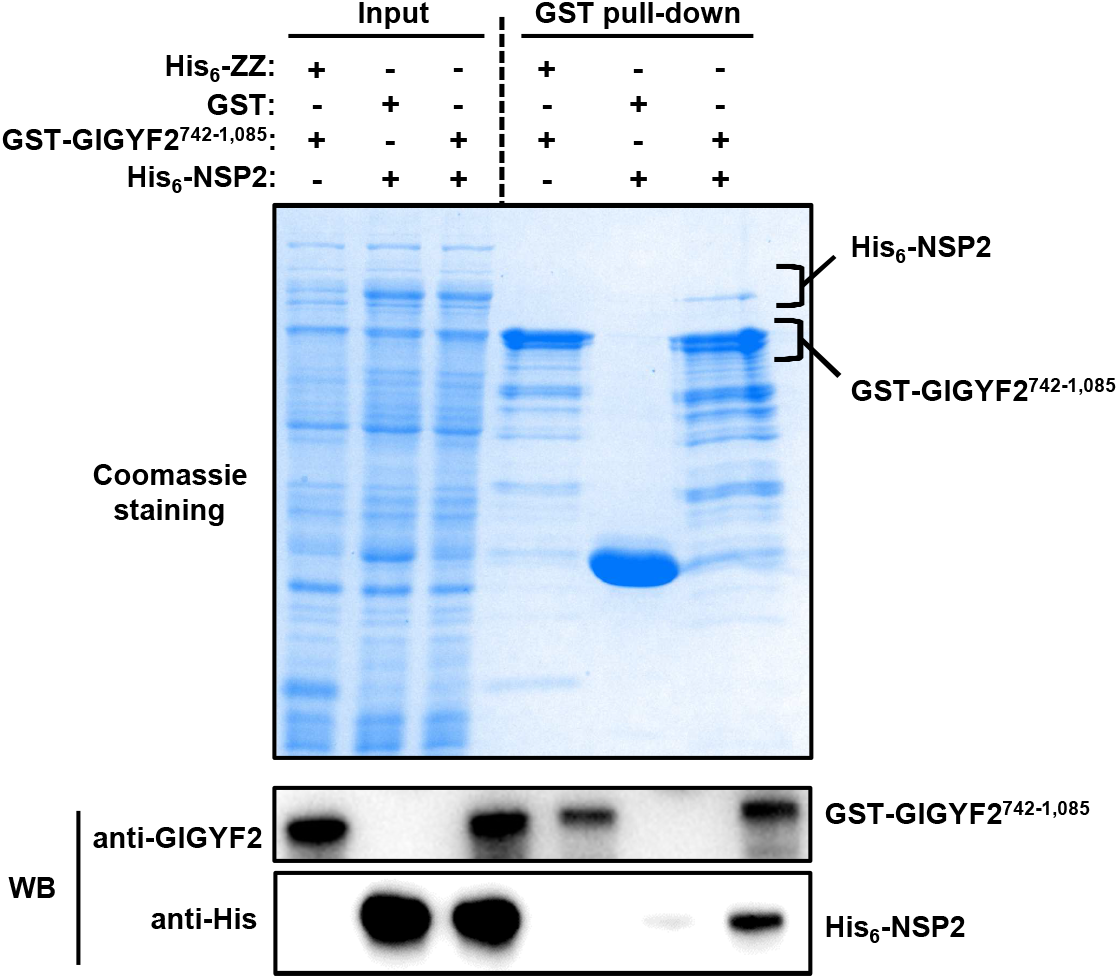
GST pull-down assay showing the interactions between recombinant GST-fused GIGYF2^742-1,085^ and His_6_-NSP2. Both His_6_-NSP2 and GST-GIGYF2^742-1,085^ were co-expressed in *E.coli* and GST pull-down assays were performed using an *E. coli* lysate. The starting material (Input) and bound fractions were analyzed by SDS-PAGE followed by Coomassie blue staining and Western blot with the indicated antibodies. GST served as negative control.

**Supplementary Figure 3.**
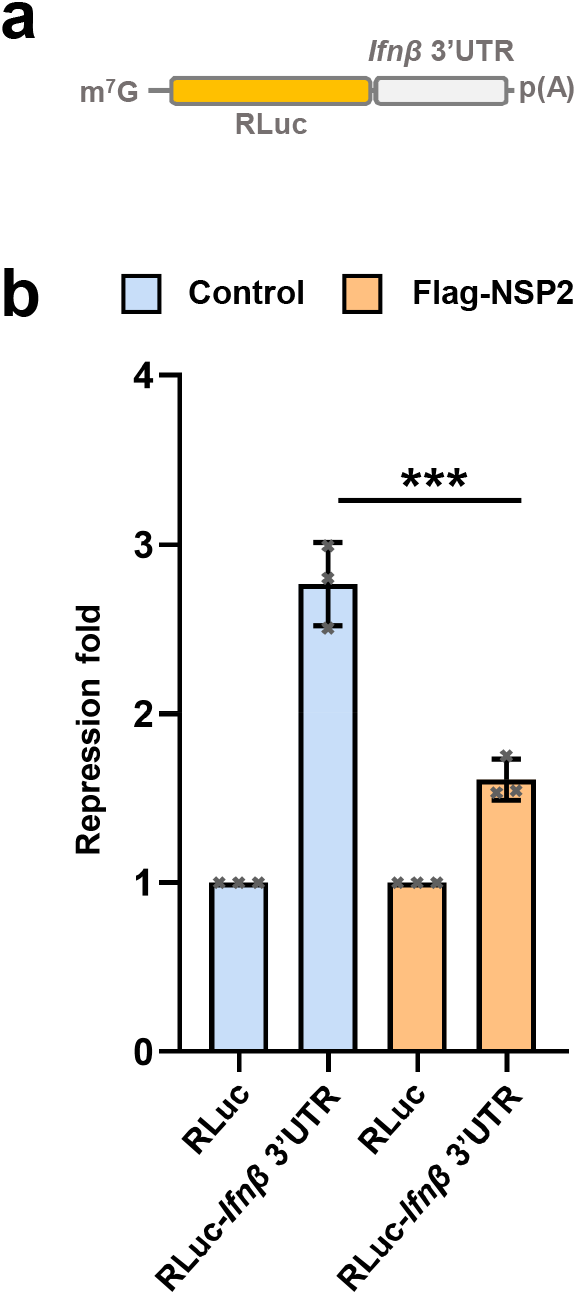
**(A)**Schematic of the psiCHECK2-RLuc-*Ifnb1* 3’ UTR reporter. **(B)**HEK293T cells were co-transfected with psiCHECK2-RLuc-*Ifnb1* 3’ UTR reporter or the psiCHECK2 reporter (as control). Vectors encoding Flag-NSP2 or Flag (empty vector) were also added in the transfection mixture. Luciferase activity was measured 24 h after transfection. RLuc values were normalized against FLuc levels, and the repression fold was calculated for the psiCHECK2-RLuc-*Ifnb1* 3’ UTR reporter relative to the psiCHECK2 reporter level for each condition. Data are presented as mean ± SD (n = 3). (***) *P* < 0.001 (2-tailed Student’s t test).

